# AngoraPy: A Python Toolkit for Modelling Anthropomorphic Goal-Driven Sensorimotor Systems

**DOI:** 10.1101/2023.10.05.560998

**Authors:** Tonio Weidler, Rainer Goebel, Mario Senden

**Affiliations:** Department of Cognitive Neuroscience, Faculty of Psychology and Neuroscience, Maastricht University, 6200 MD, Maastricht, The Netherlands

**Keywords:** goal-driven, modeling, deep learning, reinforcement learning, sensorimotor, toolbox, anthropomorphic robotics, recurrent convolutional neural networks

## Abstract

Goal-driven deep learning is increasingly used to supplement classical modeling approaches in computational neuroscience. The strength of deep neural networks lies in their ability to autonomously learn the connectivity required to solve complex and ecologically valid tasks, obviating the need for hand-engineered or hypothesis-driven connectivity patterns. Consequently, goal-driven models can generate hypotheses about the neurocomputations underlying cortical processing. Whereas goal-driven modeling is becoming increasingly common in perception neuroscience, its application to sensorimotor control is currently hampered by the complexity of the methods required to train models comprising the closed sensation-action loop. To mitigate this hurdle, we introduce *AngoraPy*, a modeling library that provides researchers with the tools to train complex recurrent convolutional neural networks that model sensorimotor systems.

## 1 Introduction

Goal-driven modeling is a novel approach in computational neuroscience that utilizes deep learning to construct highly performant models of brain function (Yamins and DiCarlo, 2016). By relying on computational optimization, this approach obviates the need for hypothesis-driven construction and hand engineering of brain models. Instead, such models emerge when deep neural networks with an appropriate architecture (e.g., layers resembling interareal pathways and topography), appropriate activation functions, and biologically meaningful input modalities are trained on natural stimuli and ecologically valid tasks. The resulting models can be probed *in silico* to yield hypotheses about yet-undiscovered computational mechanisms in the targeted structures. These, in turn, can then be tested *in vivo*. The enormous potential of this approach has been attested by recent successes within the neuroscience of perception (Yamins and DiCarlo, 2016). In particular, convolutional neural networks (CNNs) proved to be effective models of visual (e.g., Kubilius et al., 2018; Schrimpf and Kubilius, 2018; Yamins et al., 2014) and auditory (e.g., Li et al., 2022; Kell et al., 2018) cortex. Linear neural response prediction (Yamins et al., 2014; Cadieu et al., 2007; Carandini et al., 2005; Yamins and DiCarlo, 2016), representational similarity analysis (Kriegeskorte and Douglas, 2018), or composite metrics like brain-score (Schrimpf and Kubilius, 2018) have repeatedly validated learned representations and their hierarchy as at least reminiscent of representations in the brain.

However, at present the advantages of goal-driven deep learning are limited to the study of perception and do not translate well to the study of the sensorimotor system. At the same time, this domain would have much to gain from the goal-driven approach. The limitations of hand-engineering already apparent when modeling open loop systems such as sensory cortices become even more prominent for the closed perception-action loop: The complexity of the sensorimotor system renders construction of a complete computational account by hand untenable (Loeb and Tsianos, 2015). Moreover, reductionist models limiting themselves to an isolated phenomenon neglect the idea of emergence (Ellis, 2008) which postulates that unique properties or behaviors of a system also arise from the *interactions between* its components. Yet, hardly any research has thus far been conducted in the direction of goal-driven sensorimotor models. This is because goal-driven deep learning is typically based on data-intensive supervised learning. While datasets required to optimize models to perform perceptual tasks are widely available, those necessary to optimize models on sensorimotor tasks are sparse. Michaels et al. (2020) collected recordings from tracking gloves worn by monkeys during a grasping task and trained a recurrent neural network (RNN) on latent representations of simulated visual input to predict recorded muscle velocities. The goal-driven model reproduced the neural dynamics recorded in a neural interface in the monkeys’ AIP, F5, and M1. However, reproducing muscle velocities is a much narrower task than generating overt behavior. Accordingly, mapping specific input stimuli to specific muscle (or joint) activations limited to an individual’s grasping strategies might not actually represent an ecologically valid task setting. Additionally, gathering such recordings *in vivo* is costly and time-consuming. A better solution would avoid the need for labeled data. Reinforcement learning (RL) presents the (currently) most potent solution of that sort. Here, an artificial agent (e.g., a brain model) learns to adopt some behavior by autonomously exploring a task environment and adapting its decision making to maximize an external reward that indicates the task objective. Recent advances in deep learning-based robotic control (OpenAI et al., 2020, 2019; Huang et al., 2021) demonstrate the ability of this approach to teach RNNs complex, ecologically-valid tasks such as in-hand object manipulation *in silico*. Unfortunately, training such systems requires substantially more computation and tuning than classification tasks, and applying RL algorithms to recurrent convolutional neural networks (RCNN, where *convolutional* usually describes the network’s ability to process visual information) is a non-trivial engineering effort. Therefore, we believe that goal-driven sensorimotor control research would dramatically benefit from a flexible but easy-to-use toolkit for training complex biologically constrained neural networks in an efficient RL setup.

To address this, we introduce *AngoraPy* (**An**thropomorphic **Go**al-directed **R**esponsive **A**gents in **Py**thon), a python library for the efficient, distributed training of large-scale RCNNs on anthropomorphic robotic tasks with human sensory input modalities. AngoraPy is explicitly tailored toward the use of goal-driven deep learning within computational neuroscience. It bridges the technical gap between modeling deep neural network representations of the brain and training them on complex tasks at scale. It thereby facilitates the use of goal-driven deep learning for sensorimotor research by requiring no profound knowledge about the underlying machine learning techniques from its user. In the following, we first give a brief overview of modeling the sensorimotor system and reinforcement learning (Section 2). Subsequently, Section 3 describes the framework in detail alongside an exemplary use case. AngoraPy aims to provide a flexible tool that is agnostic to task and model definitions. To showcase this flexibility, we present both the results of the exemplary case study and a battery of benchmarks that demonstrate out-of-the-box performance on diverse task sets and networks (Section 4). We then conclude by summarizing our work and discussing prospects for future applications (Section 5).

## 2 Background

Research on sensorimotor control investigates the mapping of sensory stimuli to motor commands constituted by the coupling between sensory cortices and motor cortex. Taking into account the effect of motor action on sensory stimuli (both directly and indirectly), this coupling establishes a closed-loop feedback control system. The study of its controller, the nervous system, often starts with a hypothesis embedded in or abstracted from a model. For instance, trajectory control was an influential model of movements under perturbation but was later shown to be incomplete when the goal is not the trajectory itself but a static target (Cluff and Scott, 2015). Models of metabolic muscular energy consumption often base their cost functions on the heat and power observed in individual or groups of muscles (Tsianos et al., 2012). Other work uses kinematic measurements and electromyography (EMG) data to select the most suitable amongst various candidate minimization criteria (Pedotti et al., 1978). All these models begin with a hypothesis about a mechanism sourced inductively from behavioral or physiological data.

Another line of research is concerned with the functional roles of the different cortical areas involved in sensorimotor control. To this end, decoding studies using, e.g., functional magnetic resonance imaging (fMRI) have successfully identified neural representations related to movement parameters. For instance, Mizuguchi et al. (2014) have shown that right frontoparietal activity reflects the intended force level in grasping. Using multi-voxel pattern analysis in fMRI data, Gallivan et al. (2013) found activity in frontoparietal regions that predicts the movements of the participating limbs and the actions of the hand during grasping. Similarly, Filimon et al. (2015) discovered that the voxels that encode action-related information during reaching movements spread over vast portions of both the premotor and posterior parietal regions. These lines of work started with a hypothesis (or several) about a candidate role for some region, pathway, or network. However, research that simultaneously elucidates the operation of large sensorimotor networks and the contribution of individual regions remains scarce (Gallivan et al., 2018).

Although valuable in their own right, none of the current approaches can provide a comprehensive perspective on the sensorimotor system. First, hand-engineered models scale poorly to more intricate processes involving larger cortical networks. Second, from the perspective of Marr’s three levels of analysis (Marr, 1982), these models only touch on the first two levels of explanation. The implementation of the mechanisms (i.e., the third level) in terms of neural circuits and biological constraints still requires substantial work (Franklin and Wolpert, 2011). This is also and maybe particularly true regarding the study of neural transformations, rendering even the second level only partially covered. While decoding studies provide valuable information about the information encoded in various regions, they cannot provide insights into the neurocomputational strategies by which these representations are transformed within and across brain regions. Due to the complexity of these transformations, it is also near impossible to hand-engineer them. Similarly, it is unlikely that fitting deep models to reproduce recorded neural responses in multiple areas would yield great success given that data for such an approach is lacking (Yamins and DiCarlo, 2016).

### 2.1 Reinforcement Learning-Based Goal-Driven Modeling

Reinforcement learning-based goal-driven modeling overcomes these issues. Navigating the complexity of the neurocomputations to be modeled is handled by optimization and training data is generated autonomously on-the-fly. Reinforcement learning (RL) optimizes models to approximate a mapping *π*(*s*_*t*_) → *a*_*t*_ from state descriptions *s*_*t*_ ∈ *S* to actions *a*_*t*_ ∈ *A* that maximizes the sum of rewards *r*_*t*_ allocated to a trajectory (episode) of state-action pairs. In contrast to supervised and unsupervised learning, RL does not rely on pre-collected data but on self-generated samples gathered by acting in the environment. The mapping *π*(*s*_*t*_) constitutes a (behavioral) policy parameterizable by any function approximator, but in the setting described here, the policy is implemented by a deep neural network that models (parts of) the brain. Physically (albeit in simulation), the agent is situated in its environment such that its actions affect the environment’s progression through time.

Figure 1 depicts the relation between agent and simulation, and the interaction of *body, brain* and the *environment*. Generally, this interaction follows the following schema. The environment triggers sensory stimulation in the agent’s body, which then feedbacks the vectorized perception of that stimulation to the agent’s brain. The brain maps the sensory information to a desired motor command and projects it to the body. The body executes the motor command and thereby affects the environment. The new state of the environment produces the next sensory stimulation of the body, so the circle closes. Alongside describing the state, the environment also rewards the action performed in the previous state. On the basis of this information, the brain’s parameters can be optimized.

**Figure 1.**
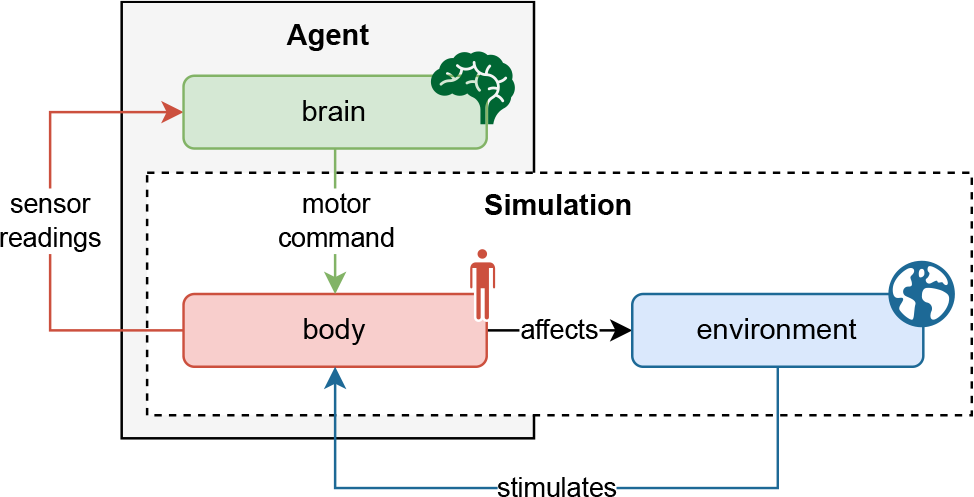
Schematic interaction between agent and physics simulation. The agent consists of a brain and a body, whereas the physics simulation covers the environment and body. As such, the agent’s body constitutes the interface between the brain and the environment. At every timestep in the simulation, the environment causes an effect on the body which generates sensory stimulation. The body’s sensors read this stimulation and communicate the information to the agent’s brain. The brain then maps the description of the perceived state to a motor command which it sends back to the body. The body executes the action and thereby affects the environment as well as itself. With readings of the new sensory state, this cycle recurs until the environment ends the episode, a trajectory of state transitions.

### 2.2 Plausibility, Validation and Hypothesis Generation

Notably, the process by which learning occurs in this setup need not be biologically plausible. It suffices that biological and artificial agents have approximately the same objective, defined by some reward-based reinforcement signal. This is an innate characteristic of any RL algorithm. Freeing the training procedure from biological constraints allows highly complex tasks to be learned in a feasible time using state-of-the-art machine learning techniques. We describe AngoraPy’s implementation of these techniques in Section 3.

Trained models can be validated with the help of, e.g., decoding studies or representational similarity analysis (Kriegeskorte et al., 2008). From validated models reproducing real cortical representations, hypotheses about, e.g., transformations can be extracted. These can then be tested by making predictions for new data on how some signal develops throughout the cortex. Therefore, goal-driven deep reinforcement learning complements hypothesis-driven research on the sensorimotor system as a method of *hypothesis generation*.

Interestingly, hypotheses can also be generated where the computational model and the experimental data do *not* fit (Loeb and Tsianos, 2015). Conditions under which such errors occur can inspire empirical research to identify relevant biological or physical constraints whose inclusion may refine the goal-driven model. *In silico* ablation studies on these constraints then shed light on their potential functional roles.

## 3 Method

In this section, we will provide an overview of the design of AngoraPy, which trains models that combine an anthropomorphic motor plant with a deep neural network. We begin by outlining our core design principles, followed by an introduction to the framework’s main dependencies. Next, we provide a technical overview of AngoraPy’s key components, including a prototypical workflow that illustrates the process of building a model of in-hand object manipulation. It is not our aim to hereby present a definitive model, but to demonstrate how researchers can utilize AngoraPy to define and train their own models.

Table 1 summarizes deployment options for AngoraPy and their intended usage. **AngoraPy is open source and available on GitHub**^1^ and licensed under GNU GPL-3.0. The repository also gives installation instructions for the different deployment options. In addition to the Python API, command-line scripts are available as entry points with extensive customization options. The software is tested and maintained under Linux, but the Python API is not specific to an operating system by design. We also maintain a growing collection of hands-on tutorials^2^ and documentation.

**Table 1:** Deployment options available for AngoraPy. Additional details and installation instructions are available on GitHub^1^.

|  | Description | Intended Usage |
| --- | --- | --- |
| **Source** | The open-source code is available on GitHub. On the repository, a master branch corresponds to the latest release version and the development branch features more recent changes and additions. | Contribution, Development Versions |
| **PyPI** | All release versions are available on the Python Packaging Index (PyPI) and can thus be easily installed via pip. | Local installation |
| **Docker** | A Dockerfile is available on GitHub and can serve as is for a clean installation of AngoraPy with all its dependencies, GPU support, and MPI support. This also allows for easy deployment on HPC clusters. | Local Installation, HPC Installation |

### 3.1 Principles and Goals of Design

We have built AngoraPy with appropriate design principles in mind: *Neuroscience First, Modularity* and *Pragmatism*. These guide the implementation toward the overall goal of providing a flexible but effective tool to neuroscientists, and draw inspiration from the principles used to build PyTorch (Paszke et al., 2019).

As we put **neuroscience first**, AngoraPy by design addresses the needs of neuroscientists who want to build goal-driven models with ease. It is our main objective to provide an intuitive API that interfaces background processes without requiring elaborate knowledge about them. This entails limiting options to only those that truly benefit neuroscientific research. AngoraPy does not aim to be a comprehensive reinforcement learning or deep learning library. In the remainder of this section, we will motivate those choices and options.

AngoraPy is built for **modularity**, where options matter. The API is general enough to support a wide range of applications. In particular, AngoraPy is not specific to tasks or models. However, to guarantee easy interaction with the framework, both need to follow requirements (we lay them out in Section 3.3 and Section 3.6). As development continues, these requirements are monitored and adjusted when needed.

Lastly, the framework is built with **pragmatism**. Computational efficiency is a matter of high importance in AngoraPy. At the same time, performance matters, and we aim to provide tools that achieve state-of-the-art results. However, neither efficiency nor performance should substantially hinder the other, nor should the simplicity of the API suffer from either. As such, we try to maintain a somewhat Pareto optimal balance between the three objectives.

### 3.2 Main Dependencies

Before detailing the framework itself, it is useful to discuss the choice of underlying libraries that provide its backbone. This choice is not arbitrary, as it has implications for the modeling supported by *users* of AngoraPy.

The rise of deep learning as a toolbox of algorithms in many fields was greatly accelerated by the increasing availability of well-designed, easy-to-use, and flexible autodifferentiation packages. Since the library introduced here trains deep neural networks as models of the brain, it also builds on such software. The two most prominent packages currently available are *PyTorch* (Paszke et al., 2019) and *TensorFlow* (Abadi et al., 2016). However, *JAX* (Bradbury et al., 2018) is also becoming popular among fundamental researchers in deep learning due to its low-level flexibility. Choosing among these options is not always obvious since each of them has sufficient functionality to support any research. In AngoraPy, our aim was to cater not only to the contributers’ needs and expertise but, of course, also to that of the end users. This is relevant since we neither aim nor consider it useful to hide the implementation of model architectures behind a native API. Instead, models are built through the API provided by the autodifferentation library. With dependence on TensorFlow, we chose a long-established option to match the experience of many researchers. TensorFlow also comes with integrated support for its high-level extension Keras (Chollet and others, 2015), providing an easy modeling interface with high flexibility. As of 2021, the combination of TensorFlow and Keras remains the most popular on StackOverflow, Kaggle, PyPi, and Google Colab (Team Keras, 2021).

With TensorFlow+Keras, AngoraPy covers the *brain* component of the agent. To cover *body* and environment, we use a native MuJoCo+Gym stack. MuJoCo (Todorov et al., 2012) is a recently open-sourced physics simulator offering high computational accuracy and speed. Gym (Brockman et al., 2016), on the other hand, offers an interface for easy communication with an environment in reinforcement learning applications. For AngoraPy, we have built a native wrapper for Gym environments, extending their functionality to suit the needs in an anthropomorphic setting better and to simulate body parts and their interaction with the world in MuJoCo.

### 3.3 Tasks & Simulation

In goal-driven modeling, the task definition is crucial to the methodology’s premises. It sets the *goal* and thereby *drives* learning to generate the neurocomputations one seeks to discover. However, prescribing appropriate task and simulation constraints is not trivial.

AngoraPy ships an API specifically designed for anthropomorphic tasks. It extends the Gym library (Brockman et al., 2016) by interfaces and wrappers that inject additional features and standards tailored towards sensorimotor applications. This comprises adapted interfaces to the official Python bindings of MuJoCo (Todorov et al., 2012) and for implementing anthropomorphic task environments. Several built-in dexterity tasks employing a simulation of an anthropomorphic robotic hand (the Shadow Dexterous Hand^3^) exemplify such implementations and can serve as a foundation for dexterity research. They are implemented on top of a hand model and task implementation originally included but currently discontinued in Gym.

**Example**. *In the current and following example segments, we will exemplify the construction of a goal-driven in-hand object manipulation model in AngoraPy. To this end, example segments annotate the upcoming technical descriptions with practical illustrations. In-hand object manipulation (IHOM) is a manual dexterity task. To simulate it, we use the hand model shipped with AngoraPy. It consists of 24 joints and has its palm connected to a fixed socket via a joint with two degrees of freedom. Actuators are directly attached to the joints and apply control in terms of the absolute desired joint angles. Of the 24 joints, 4 are coupled. Thus, they cannot be controlled directly but move dependent on other joints. Accordingly, the motor plant has a total of 20 degrees of freedom. In-hand object manipulation covers a broad category of tasks, but teaching it to an artificial agent requires a prototypical specification. Consistent with OpenAI et al. (2020), we prototype the manipulation task as the in-hand reorientation of a cube whose faces are uniquely colored and labeled. A target reorientation is specified as an angle of rotation around a fixed point (the object’s center) and achieved if the cube’s rotation angle lies within η units of the target angle; that is, their distance d_g_(t) ≤ η. To encourage stable behavior towards the end of a reorientation, we define a single episode as a chain of reorientations. Thus, the agent needs to learn manipulation in a manner that maintains sufficient control to enable it to perform the next reorientation from the endpoint of the previous. Per reorientation (i.e., goal), the agent is given 8 seconds and the total number of possible reorientations is capped at 50. The 8s-timer resets for every reached goal. The episode ends immediately when the cube is dropped, indicated by the cube center’s z position coming below that of the palm*.

When AngoraPy instantiates an environment, it encapsulates it with a wrapper connecting external (native Gym) environments with its own API. Specifically, state descriptions are converted to *Sensations*, the canonical input type for AngoraPy, which we will introduce in Section 3.4. Optionally, the wrapper accepts transformers modulating observed states and rewards. AngoraPy offers two built-in transformers which normalize rewards and states, respectively, via running mean estimators. Such transformations can be crucial in many tasks (Ilyas et al., 2020) but hamper progress in others. For instance, reward normalization is not useful with environments assigning constant rewards for nonterminal states. As the number of steps per episode approaches infinity, the reward for non-terminal states approaches zero, while that for terminal states approaches (negative) infinity. As episodes last longer, the relative importance of the terminal state thus increases, which potentially destabilizes learning.

A key component of an RL environment is furthermore the reward function. In AngoraPy, reward functions integrate into environments as the union of a *reward prototype function* and a *reward configuration*. This setup enables parametrization and inheritance and thereby facilitates objective-related ablation experiments. Additionally, non-static reward functions allow for shifts in the balance between different terms in the reward function. These shifts can create training schedules that facilitate learning.

**Example**. *To model the IHOM objective via a reward function, we provide feedback on success (r*_*s*_*), failure (r*_*f*_ *), and progress (r*_*p*_),

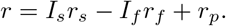

*The indicator variables I*_*s*_ *and I*_*f*_ *are equal to* 1 *on success and failure (dropping), respectively (and* 0 *otherwise). Progress r*_*p*_(*t*) *at time t is defined as*

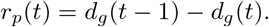

*The progress term r*_*p*_(*t*) *also captures regression. During training, a reward transformer estimates the running mean and variance of the reward function to normalize its output*.

### 3.4 Sensation

To learn the task, the agent must find a favorable mapping between states and actions. AngoraPy trains biologically inspired networks to control anthropomorphic, simulated bodies. Accordingly, the state *s*_*t*_ on which the policy *π*(*s*_*t*_) conditions its action distribution needs to emulate human sensory modalities. At present, these modalities comprise *vision, touch*, and *proprioception*, as they are key components of human sensorimotor processing (Wolpert et al., 2011). Although the task goal could also be implicitly encoded in either of these, states additionally feature an explicit *goal* description. One may consider these explicit goal representations internal representations, as encoded in, e.g., areas of the prefrontal cortex (Miller and Cohen, 2001; Braver, 2012). Together, this 4-tuple constitutes the (vectorized) input to the model, contained within the Sensation type. Keeping the tuple’s elements separate allows for multiinput architectures, as required when modeling sensory modalities that exclusively target their respective sensory cortices.

**Example**. *For manipulation, all three sensory modalities are relevant feedback. However, to simplify the problem for the purpose of this example, we omit vision and replace it with immediate information on the object’s pose. Additionally, the model requires a goal description. The environment returns this as a vector representation of the target quaternion. A transformer (as described in Section 3.3) records the means and variances across all modalities and uses these to normalize the states*.

#### 3.4.1 Auxiliary Data for Asymmetric Value Functions

Importantly, goal-directed modeling aims at plausible inference. As discussed in Section 2, the learning procedure itself, however, need not be biologically plausible for its premises to hold. During training, this distinction can be leveraged. First, this concerns the learning algorithm that can be applied. Second, we can enrich the state with auxiliary information only used during training (Pinto et al., 2018). The training algorithm used by AngoraPy requires a value network *V* ^*π*^(*s*_*t*_) that estimates the expected value of being in state *s*_*t*_ given the current policy *π*. Since *V* ^*π*^ is discarded at inference time, it can rely on any non-biologically plausible, *auxiliary* input without harming the model’s plausibility.

**Example**. *We provide auxiliary information to the value network during training. These include fingertip positions, the relative orientation between the target and the current object orientation, the object’s positional velocity, and the object’s rotational velocity. Together with sensory and goal inputs, these constitute the Sensation instances produced by the manipulation environment. AngoraPy automatically feeds these to the relevant parts of the model*.

### 3.5 Motor Commands

An agent’s brain maps its sensations to motor commands. AngoraPy trains stochastic policies and thus maps to the parameters of probability distributions. The executed motor commands are directly sampled from that distribution. Like its human counterpart, an anthropomorphic agent has continuous control over multiple action dimensions. This entails the approximation of a *joint* probability density function (PDF). One assumes the joint distribution over possible actions to be i.i.d. marginal distributions. In practice, modeling joint PDFs can become problematic because the space of potential actions is infinite. With a growing number of marginal distributions, this makes predictions about interactions difficult. Therefore, it proves beneficial to bin each continuous action variable in applications with many DoF (OpenAI et al., 2020, 2019). This converts the problem to the approximation of a multicategorical distribution.

AngoraPy offers different options for modeling the probability distribution of a multiple-DOF continuous policy: the multicategorical (binned), Gaussian, and Beta distributions and an interface for implementing custom policy distributions. In the following sections, the three built-in options are outlined alongside their (dis)advantages. Figure 2 depicts their characteristic shapes.

**Figure 2.**
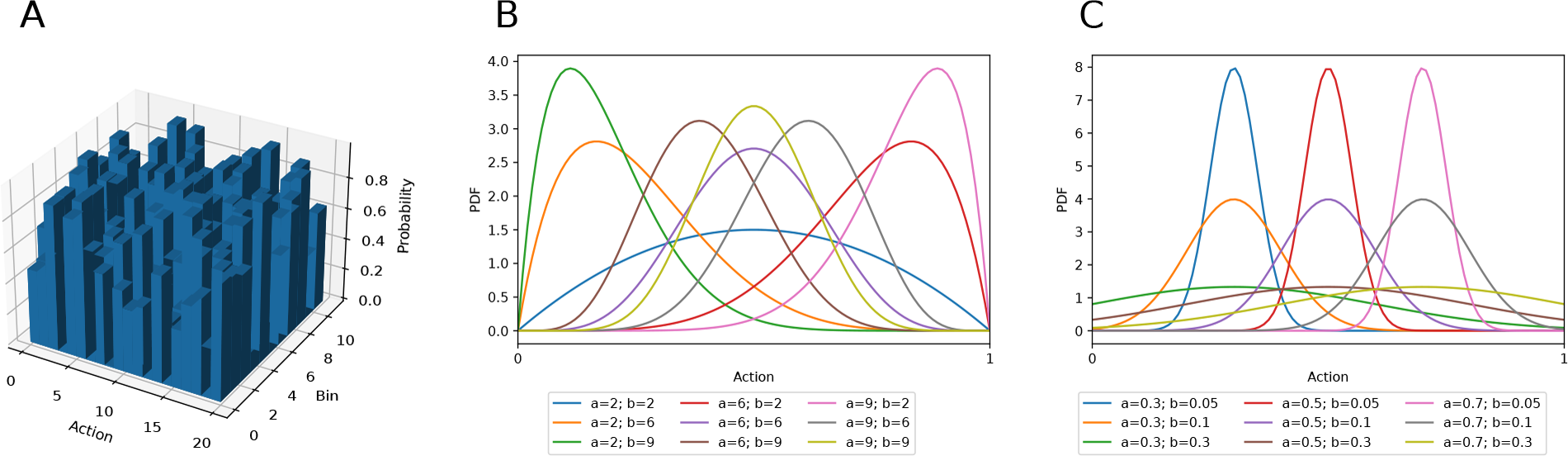
Three different policy distribution classes as described in Section 3.5. **(A)** Multicategorical distribution. **(B)** Beta distribution for different combinations of *α*- and *β*-values. **(C)** Gaussian distributions with different means and standard deviations. For equal *α* and *β* parameters, the Beta distribution resembles the Gaussian distribution. For diverging parameters, it becomes increasingly skewed. Importantly, the Beta distribution’s domain is entirely confined to the interval [0, 1].

#### 3.5.1 Multicategorical Control for Continuous Action Spaces

In the multicategorical setting, a continuous action space *A* consisting of *n* actions *a ∈* [*a*_*min*_, *a*_*max*_] is segmented into *b* bins per action, each spanning 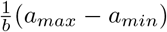 units of the continuous space. Accordingly, the model has an output space *A* ∈ ℝ^*b×n*^. Compared to a continuous policy distribution, this tends to be substantially larger. However, the problem is simplified for two reasons: First, binning the action space leads to a finite set of possible actions that the robot can execute. In contrast, a theoretically infinite number of actions can be sampled from the continuous distribution. Second, over a finite set of possible actions, optimization has full control. A multicategorical distribution can have any shape imaginable, while Gaussian and Beta distributions have characteristic shapes (Figure 2).

#### 3.5.2 Gaussian-Distributed Continuous Control

Gaussian policy distributions predict the means and standard deviations of every marginal distribution. The output space of an *n*-DoF control task is thus 2*n*-dimensional. Standard deviations are usually better trained independent from the input. That is, the standard deviations are themselves a set of directly optimized parameters, but the input state is irrelevant to their manifestation in the policy output. Essentially, the optimization then slowly anneals the standard deviations to 0 as it becomes more confident in predicting the means. Predicting the standard deviation as a function of the input is possible but less stable when facing outliers in the state space. It also adds additional load to the optimization because it requires approximating an evolving mapping between instances and confidence.

#### 3.5.3 Beta-Distributed Continuous Control

In bounded action spaces, the infinite support of Gaussian distributed policies biases them towards the limits (Chou et al., 2017). The unconstrained support requires a fold of all predictions outside of an action’s bounds onto the boundary values and artificially increases their probability. Chou et al. (2017) demonstrated that an agent predicting the parameters of a Beta distribution circumvents this issue. The support of the Beta distribution is in the interval [0, 1], independent of the parameters, and is parametrized by *α* and *β* (Figure 2). By scaling samples to the allowed interval of the action, the prediction is complete but bias-free. Chou et al. (2017) showcased the positive effect on TRPO (Schulman et al., 2015) and ACER (Wang et al., 2017) agents, but we also found this to be useful in PPO (see Section 4.2). In motor control, actions are constrained by maximum joint flexion and extension. The Beta distribution hence is a natural choice for many anthropomorphic systems.

**Example**. *For IHOM, we train the agent with a multi-categorical distribution. The model then predicts a vector a* ∈ ℝ^*db*^ *where d is the degrees of freedom of the motor plant (*20 *for the ShadowHand) and b is the number of bins per degree of freedom. Consistent with OpenAI et al*. *(2020), we set b* = 11.

### 3.6 Brain Models

With **s**_*t*_ and **a**_*t*_ in *π*(**s**_*t*_|*θ*) → **a**_*t*_ covered in the previous sections, the following focuses on the neural network models parametrized by *θ*. We argued in Section 3.2 for TensorFlow+Keras as the backbone of their implementation. AngoraPy can train any model implemented in Keras if it adheres to two constraints. First, the input layers of the model must be a subset of the modalities present in a Sensation. Second, the model must be available as a tuple of policy, value and joint networks, where joint is the combination of policy and value networks into a single network with two heads. The policy network’s output must be built by the agent’s policy distribution and the value network must map the input onto a single scalar. These requirements are depicted in Figure 3.

**Figure 3.**
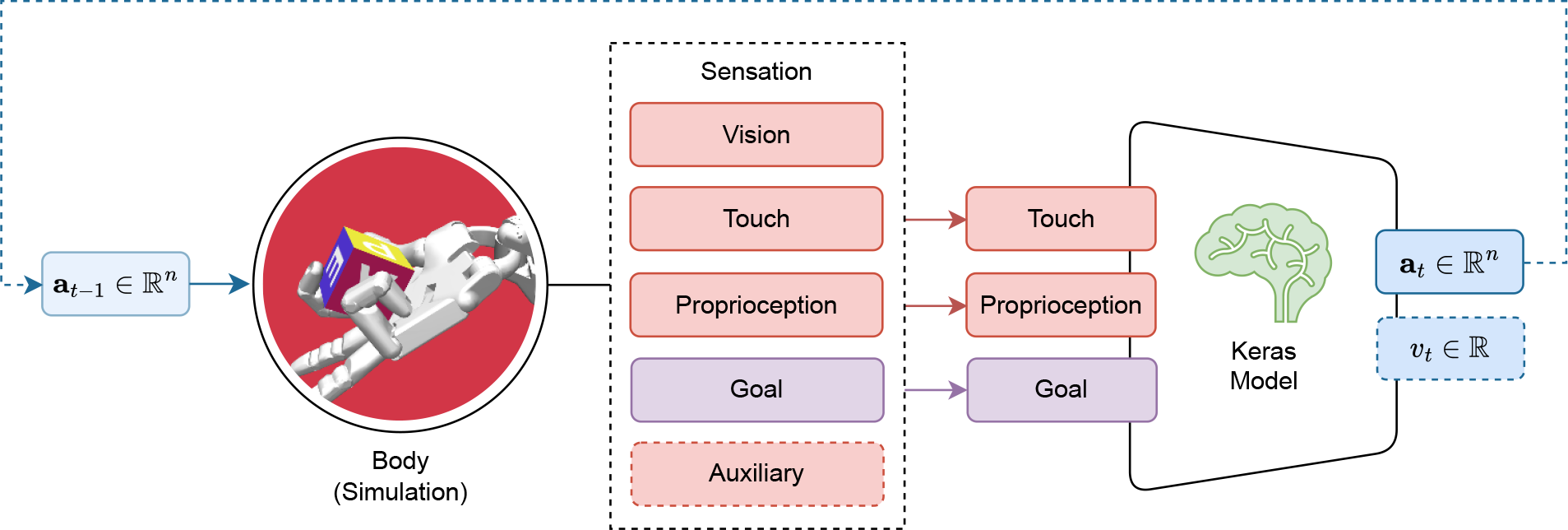
Requirements for models trained with AngoraPy. The set of input modalities must be a (sub)set of the modalities available in a Sensation. Outputs must be a policy head output following the desired distribution’s shape, followed by a value output head projecting to a single scalar.

Often, a deep model of the sensorimotor system will require a recurrent convolutional neural network (RCNN) implementation. Recurrence affords memory and representations of environmental dynamics. Convolutional networks mimic the hierarchical and retinotopic structure of the visual cortex (LeCun et al., 2015). Furthermore, their layers learn representations that resemble those found in the respective cortical areas (Yamins and DiCarlo, 2016; LeCun et al., 2015). During training, AngoraPy employs several strategies to deal with the computational load of RCNNs during training (Section 3.7). However, hardware will always constrain the viable complexity of a model.

**Example**. *The model we use for IHOM is a shallow but high-dimensional, recurrent network with an embedding layer, and implements the architecture used by OpenAI et al*. *(2020). Different modalities from the input Sensations we defined in a previous example will first be concatenated into a single vector and then fed into the embedding layer. The architecture is available in AngoraPy’s network registry (under the key wide). Recurrence is necessary to model the dynamics of the motion of external objects because neither rotational nor positional velocities are directly available in the raw sensory state space*.

#### Weight Sharing Between Policy and Value Networks

Because at train time, optimization relies on state value estimates, networks need additional *value heads*. These are trained to predict the average future reward received under the current policy when in the given state. As we explained in our discussion of asymmetric value functions (Section 3.4), the value head can be discarded at inference time. Its architecture, therefore, needs not be biologically plausible. However, when designing a network architecture, incorporating the value head plays an important role. Essentially, one must decide whether to entirely separate the value from the policy network or at what stage policy and value heads diverge. Separation can lead to better peak performance since optimization need not balance between two objectives. However, since value estimation and action selection will overlap in the information they need to extract from the input, sharing the stem of their networks can lead to faster convergence. Naturally, auxiliary inputs can only be integrated where heads diverge.

**Example**. *We build the architecture without a shared base to prioritize performance over convergence time. The policy network and the value network have the same architecture up until after the recurrent layer. The policy network culminates in a distribution head generated by the PolicyDistribution instantiated previously. The value head projects to a single linear scalar predictive of the value of the state described by the given input*.

#### Modularity and Pretraining

Sensorimotor systems are vast and include various cortices. Generally, this tends to make deep implementations of sensorimotor systems inherently modular. For instance, subnetworks that model sensory cortices are generally interchangeable. Extensive research on goal-driven models in the sensory domain allows modelers to easily exploit this modularity. First, researchers investigating specific cortical areas in the context of sensorimotor processing can enrich their model by integrating it with other models. On the other hand, different models of the same brain structure can be plugged into existing sensorimotor models to evaluate their effect on functional performance and neurocomputational validity. Finally, downstream models like sensory cortices can be pretrained on functions identified in perception research. This severely disburdens the RL training on the main task. Naturally, it might also prove beneficial for the validity of upstream neurocomputations if downstream feature extraction is bootstrapped with less task specificity.

### 3.7 Training Agents

Before, we described building models and tasks and their interaction as communicated by sensations and motor commands. This sets the rules for the core of AngoraPy, the distributed deep reinforcement learning backend. AngoraPy employs proximal policy optimization (PPO) (Schulman et al., 2017) to train its agents’ policies using stochastic gradient descent (specifically Adam; Kingma and Ba, 2017). PPO is an RL algorithm that alternates between data *gathering* and *optimizing* in continuous cycles. During the gathering phase, experience is collected by acting in the environment following the current policy. During the optimization phase, this experience is used to update the policy. This asynchronicity between data generation and optimization allows for distributed, sample-efficient training, as Section 3.7.3 will detail.

#### 3.7.1 Proximal Policy Optimization

Proximal policy optimization (Schulman et al., 2017) directly optimizes the selection of actions. This avoids the proxy of evaluating every action in a given state. Gradients are calculated w.r.t. the parameterized policy *π*_*θ*_(**a**_*t*_|**s**_*t*_). In essence, the objective tries to move the policy in a direction that increases the likelihood of beneficial actions and decreases that of disadvantageous ones, captured by advantage estimate *Â*(**a**_*t*_, **s**_*t*_) ∈ ℝ. However, traversing the policy space in this way might apply steps too large to permit stable learning. PPO addresses this issue by scaling the update by the ratio *r*_*t*_(*θ*) between the currently optimized *π*_*θ*_(**a**_*t*_ | **s**_*t*_) and the previous version of the policy at gathering time 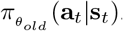. By clipping this ratio, potentially excessive updates are avoided. This is implemented in the following objective:

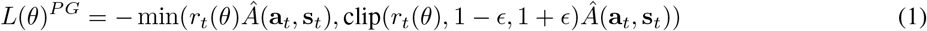

Here, 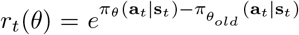 is the logarithmic ratio. The minimum in Equation 1 removes the lower bound if the advantage is positive (Schulman et al., 2017). If the advantage is negative, we want to decrease the ratio. The clipping bounds by how much we want to decrease it. The minimum ensures that recoveries from deteriorated policies receive gradients (instead of being clipped away).

The advantage *Â* (**a**_*t*_, **s**_*t*_) is the difference between the return collected from taking action **a**_*t*_ and the average future return 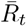 from being in state **s**_*t*_, estimated by the value network *V*_*θ*_(**s**_*t*_) with a mean squared error loss

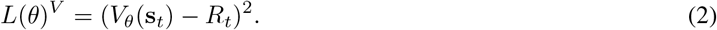

The *R*_*t*_ that serve as a target to *V*_*θ*_(**s**_*t*_) come from the same samples the policy network learns from. Thus, *V*_*θ*_(**s**_*t*_) boils down to a state-conditioned running mean estimator. The estimator may be a separate network or share part of its parameters with the policy network. In any case, it is convenient (and required for the latter) to implement the optimization of both *V*_*θ*_(**s**_*t*_) and *π*_*θ*_(**a**_*t*_|**s**_*t*_) jointly, hence combining their objectives into one joint cost function

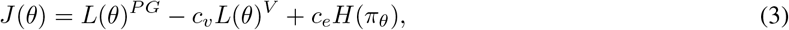

where *c*_*v*_ can be adjusted to prevent *J*(*θ*)^*V*^ from dominating *J*(*θ*)^*PG*^ when sharing parameters between value and policy network. *H*(*π*_*θ*_) is the entropy of random variable *π*_*θ*_. A stochastic policy’s entropy can be seen as a measure of the degree to which the agent still explores. Thus, incorporating it (scaled by constant *c*_*e*_) into the objective discourages premature convergence (Williams, 1992; Williams and Peng, 1991).

**Example**. *Since we previously made the decision not to share parameters between policy and value network, setting c*_*v*_ *is unnecessary. The clipping range is kept at ϵ* = 0.2, *the value recommended by Schulman et al*. *(2017). We include an entropy bonus. In the literature, the entropy coefficient c*_*e*_ *is often in the range* [0, 0.01] *(Schulman et al*., *2017*; *Engstrom et al*., *2020*; *OpenAI et al*., *2020*; *Mnih et al*., *2016). We set c*_*e*_ = 0.001 *at a fairly low value because the entropy of independent marginal distributions is the sum over marginal entropies, which naturally increases the entropy bonus*.

#### 3.7.2 Truncated Backpropagation Through Time

During training, long input sequences to recurrent layers can overload memory. That is because backpropagation through time (BPTT) needs to record the activations throughout the entire sequence to calculate gradients. Truncated BPTT (TBPTT; Williams and Peng, 1990) tackles this issue by dividing the sequence into subparts, over which the gradients are backpropagated. At the end of each subsequence, the RNN’s state(s) are passed over to the next subsequence, but the computation graph is cut off. This alleviates the need for memory in sacrifice for well-modeled long-distance dependencies (LDD) and precise gradients. However, in practice, LDDs are often negligible if we hypothesize the recurrence to encode external dynamics mostly. Imprecise gradients have likewise proven to be of limited harm. AngoraPy thus natively uses TBPTT to balance precision and memory efficiency. Note that TBPTT is often compulsory where episodes are long and networks are large. However, where this is not the case, TBPTT can be turned off by setting *s* to the length of the episode.

During the gathering phase, we collect data points in subsequences of length *s* if the policy is recurrent. Each worker initializes a buffer with all-zero matrices **B** ∈ ℝ^(*h/s*)*×s×n*^ for each collected transition information where *n* is the length of the vector representing the piece of information. The buffer matrices are progressively filled with the transitions recorded while stepping through the environment. Whenever a subsequence of length *s* is filled, it is pushed to the buffer, and the next subsequence is started. If an episode finishes before the end of a subsequence, its current state is pushed to the beginning of the buffer’s subsequence, and the worker skips *s* − (*t* mod *s*) − 1 time steps (where *t* is the current time step). Given the buffer initialization, this essentially fills the remainder of the sequence with zeros. During optimization, these zeros are masked to be ignored when calculating gradients. Tuning *s* attempts to fit the number of previous time steps relevant to modeling the dynamics of the environment. Thus, it is task-dependent. Figure 4 visualizes the implementation of TBPTT in AngoraPy.

**Figure 4.**
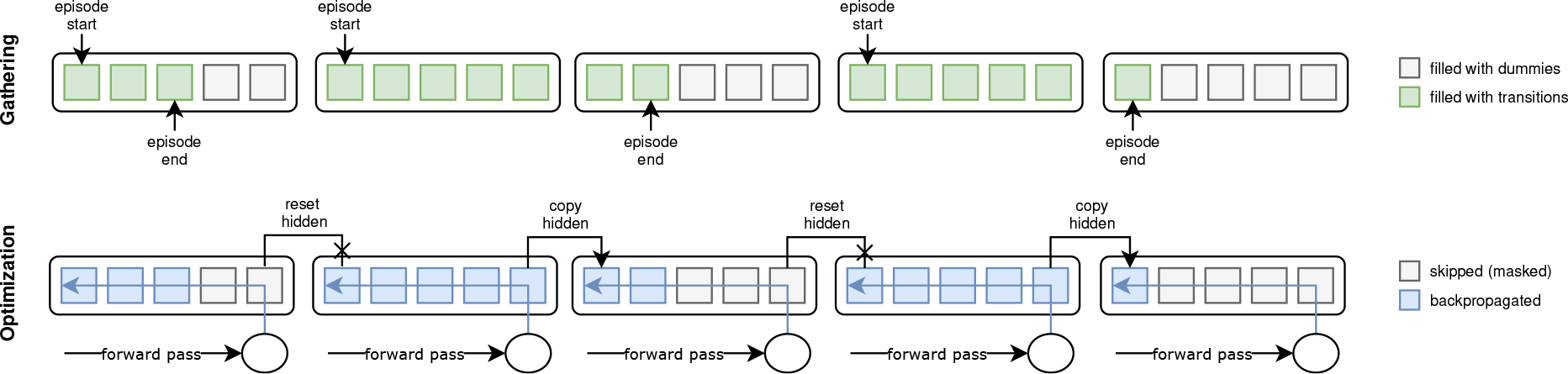
Truncated Backpropagation Through Time (TBPTT) during gathering (top) and optimization (bottom). During gathering, the transition sequence is cut off every *k* elements. If the episode ends before a cutoff point, the remainder of positions up to the next cutoff points is filled with dummy transitions. During optimization, this leads to a dataset of equally sized sequences. Dummy transitions are masked when backpropagating the error. Cutoff points at non-terminal states are handled by copying the last hidden state of the previous sequence into the initial hidden state of the current one.

**Figure 5.**
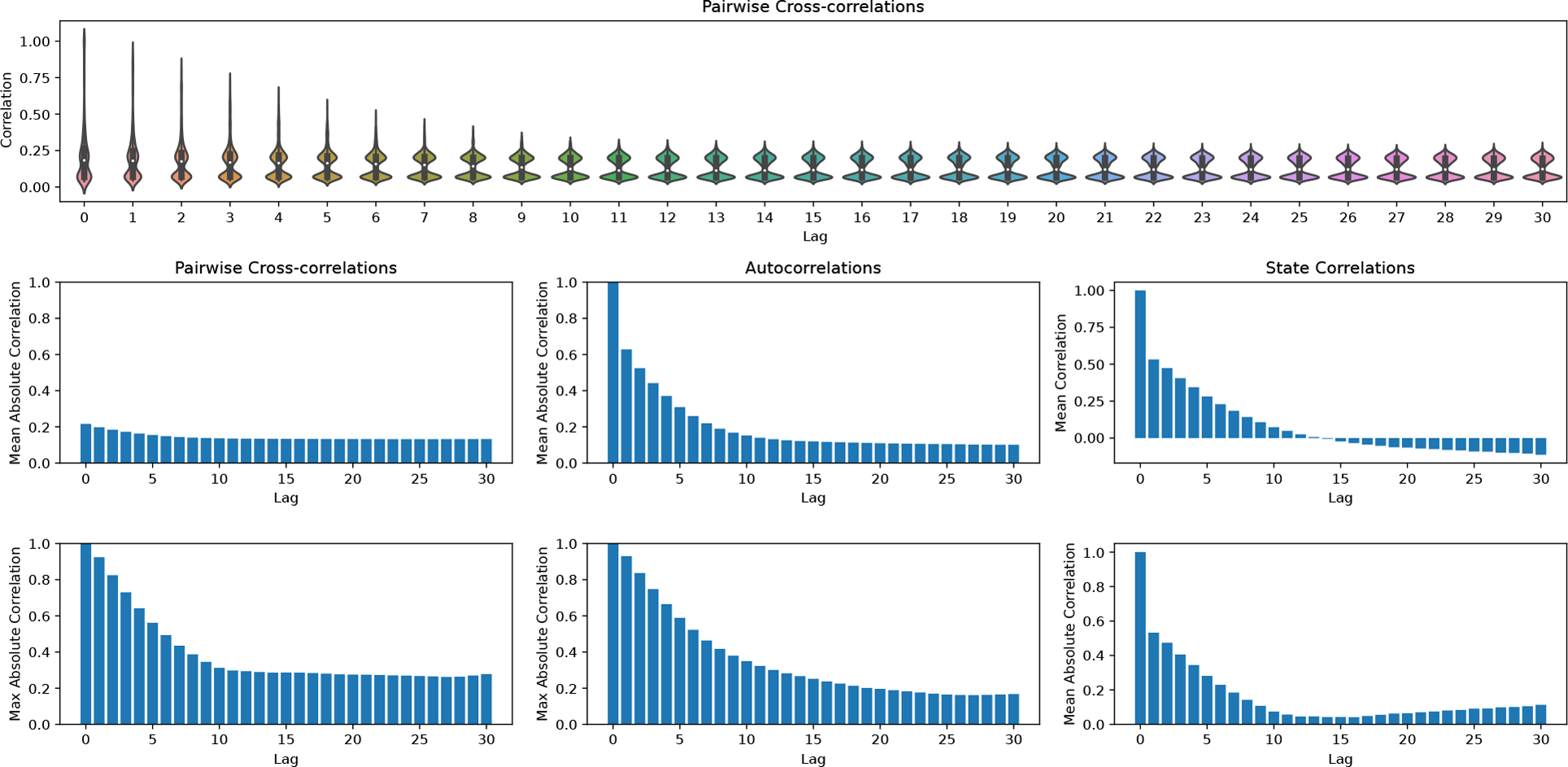
Analysis of the dynamics in the in-hand object manipulation environment to determine the temporal dependencies between state variables. (top row) Violin plots depicting the distribution of cross-correlations between singular time series at each lag. (first column) Mean and maximum cross-correlations between singular time series at different lags. (second column) Mean and maximum autocorrelations (each time series is only compared to itself) at different lags. (third column) Correlations between state vectors (as opposed to time series) taken at different lag distances.

**Example**. *To capture the dynamics of IHOM, we set s* = 16 *which equates to* 1.28 *seconds. This covers the approximation of external velocities by relating spatial to temporal distances between two consecutive time steps. As an additional means of determining a reasonable s, one can consult an analysis of the temporal dependencies of states in the environment. Consider Figure 5 where we plot different perspectives on auto- and cross-correlations between steps at different lag displacements. Although looking at the maximum cross- and autocorrelations reveals dependencies beyond a lag of* 16, *it is also evident that the strongest dependencies tend to lie within lag* 16 *and thus s* = 16.

#### 3.7.3 Distributed Computation

At the core of AngoraPy’s ability to train large-scale models on complex tasks lies its native distributed design. Learning in a recurrent, convolutional setting can require extensive training on a large amount of data. To manage this during simulation and optimization, high-performance computing on a distributed system is inevitable. Proximal policy optimization provides the algorithmic foundation necessary for AngoraPy to implement this. As PPO generates data asynchronously, any number of parallel gathering workers (GW; usually allocated to a thread/core of a CPU) can simulate the environment and roll out the policy independently. Similarly, optimization can be spread over multiple optimization workers (OW; preferentially GPU-enabled) by calculating and merging gradients on partial minibatches.

AngoraPy implements this bipartite distribution strategy under the Message Passing Interface (MPI). The detailed process is depicted in Figure 6. MPI spawns multiple processes running the same script, each building a full copy of the agent. At initiation, virtual workers are allocated evenly over processes if possible. Optimally, the allocation is bijective. Forcing uneven allocation by creating a number of virtual workers not divisible by the number of available CPU workers would substantially waste computational time. During the gathering phase, every worker acts as a GW. Each GW rolls out the policy and generates a trajectory of experience up to horizon *T*. The worker then temporarily saves this trajectory on disk. During the optimization phase, the data is evenly assigned to all OWs. Every OW reads its data share from the disk and calculates a partial gradient. In between batches, the partial gradients are accumulated among all OWs, and the total gradient is applied to the policy weights.

**Figure 6.**
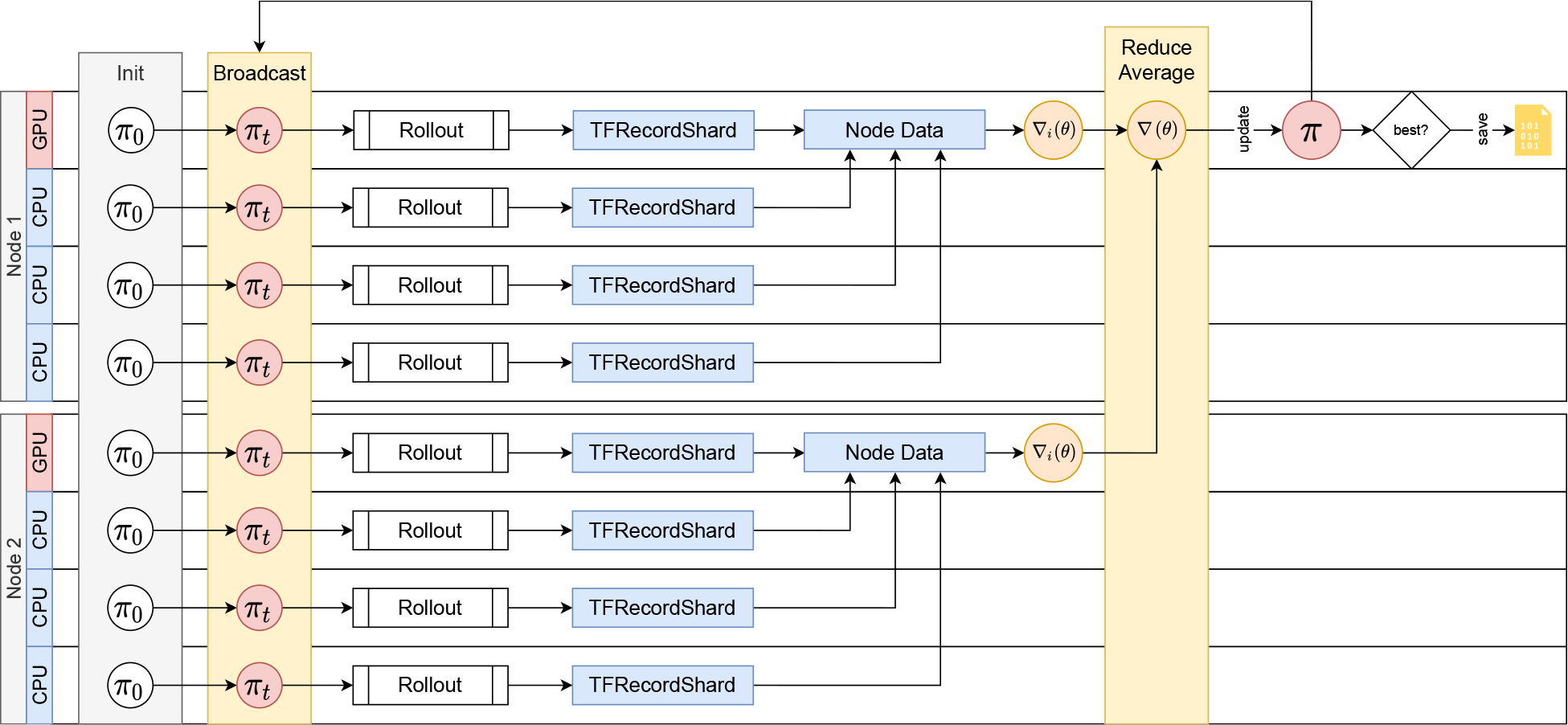
Distributed Training with MPI. An exemplary depiction of the distributed training cycle comprising gathering and optimization on two compute nodes. Distribution scales to an arbitrary number of nodes. Spawned processes (e.g., on every CPU thread or core) all share the load of gathering information. The GPU(s) on a node is assigned to the lowest-rank process of the node, and all GPU processes share the computational load of optimization. At cycle 0, every process initializes the policy, which is then synced to match the single initial policy on the root process. Every process then rolls out the policy to generate experience up to horizon *T*. The data is stored in shards. Every optimization process then collects and merges the shards produced by workers on the same node. Based on this data, the optimization process calculates a gradient. The gradients of all optimizers are then reduced into one by averaging and applied as an update to the policy on the root process. *π*_new_ is then broadcasted to all processes, which repeat the cycle by rolling out the new policy.

Note that while complex tasks involving recurrent and convolutional networks may require distributed computation, many simpler tasks such as finger oppositions or anthropomorphic robots with less degrees of freedom (e.g. locomotion) may not. AngoraPy can flexibly be applied without, with little (e.g. on local workstations or laptops) or extensive (on supercomputing clusters) parallelization. On systems without GPU support, AngoraPy instead distributes the optimization amongst CPU processes. Distributed computation is thus purely a feature, not a constraint.

##### Batch Tiling

The combination of distributed optimization and the division of sequences into chunks that need to remain in order requires batches to be tiled. Tile shapes are defined by the number of trajectories *n* and chunks *m* per trajectory where *nm* must equal the number of chunks per update. Balancing between the two presents a tradeoff between temporal bias and trajectory bias. The more chunks included per trajectory, the lower the bias of the update towards a specific temporal segment of the trajectories. At the same time, the resultant lower number of independent trajectories will inevitibly produce a bias towards specific trajectories. However, particularly during early training independent trajectories differ vastly in their course. Thus, temporal links between trajectories disappear early, and temporal biases dissolve. When AngoraPy automatically determines tile shapes based on the number of chunks per update and the number of available optimizers, it thus finds the valid shape with the highest number of trajectories included at the same time. To support this process, it is generally a good practice to choose combinations of batch and chunk sizes that produce chunks per update divisible by the number of available optimizers.

**Example**. *Gathering data is distributed over* 384 *CPU workers and we use* 32 *GPUs for optimization. Every gathering worker generates* 2048 *timesteps. We use a batch size of* 12288 *transitions. As TBPTT groups transitions into chunks of length* 16, *a total of* 768 *chunks are divided amongst optimizers. Every cycle entails* 64 *updates to the current policy each epoch and* 192 *updates per cycle. The batch size is intentionally high to lower the number of updates per cycle. Too many updates would cause the policy to move away from the data-generating policy. Additional updates would thus increasingly optimize based on off-policy data and conflict with the on-policy assumption of PPO’s objective function*.

### 3.8 Saving Agents, Resuming Experiments and Monitoring

Training automatically stores the model’s best-performing parameters between cycles, if the model’s performance during the gathering phase surpasses the previous best performance. Additionally, AngoraPy always saves the newest version of the model. Optionally, the training procedure can save the model’s state in intervals, for instance to later probe the model at different levels of progress. Model storing uses Keras’ built-in formats, but the parameters of the Adam optimizer and those needed to reload the agent object (mostly hyperparameters) are captured separately in NumPy and JSON files. This strategy allows agents to be loaded both for evaluation purposes *and* to resume training.

A *Monitor* can be connected to the training process to additionally log the training process. The monitor tracks rewards, episode lengths, losses, and transformer statistics and logs hyperparameters. Additionally, it records several statistics about training, including runtimes and memory usage. All log files are in JSON format and thus both human- and machine-readable. The progression traces are live updated during training and can be conveniently monitored during or after training using a native web application offering a filterable and searchable overview of all experiments stored on the machine and graphs visualizing stored progression traces and statistics for specific experiments or comparing multiple.

## 4 Results

In this section, we present the results of exemplary experiments to demonstrate the efficacy of AngoraPy’s RL backend. Specifically, we demonstrate its ability to handle different task categories and model architectures. The experiments were designed to highlight the versatility and robustness of the toolkit, and include benchmarks on various tasks, from classical control to robotics. The outcomes of these experiments provide compelling evidence of the effectiveness of AngoraPy in developing performant goal-driven anthropomorphic models of any shape and purpose.

### 4.1 In-hand Object Manipulation

To render the technical description of AngoraPy’s key components more intelligible, we provide a concrete example of applying the full framework using the simple model suggested by OpenAI et al. (2020). The results for this example are depicted in Figure 7. Agents reliably converge towards a strategy chaining an average of 30 goals. Convergence occurs after approximately 1, 500 cycles, corresponding to 96 hours of training in our setup employing 384 CPU cores (Intel Xeon E5-2690 v3) for data collection and 32 GPUs (NVIDIA Tesla P100 16GB) for optimization^4^. At this point, the agent has seen 1, 187, 512, 320 samples, equating 3.012 years of experience. The best performing agent converges at a similar speed, but chains approximately 40 goals.

**Figure 7.**
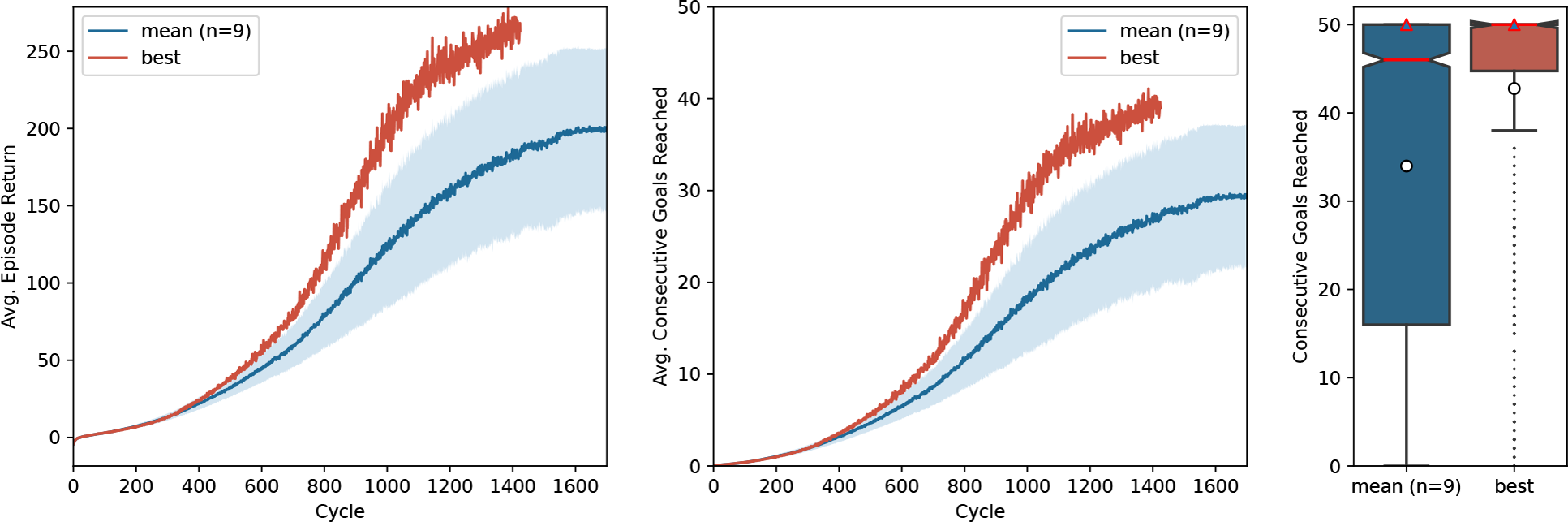
Training of in-hand object manipulation agents. Average episode returns (*left*) and average goals achieved per episode (*center*) show the progression of performance over training cycles. The mean performance over 9 independent runs is shown in blue, with a shaded 95% confidence interval. The curve of the best performing agent is shown in red. The distribution (boxplot) of 480 episodes over the number of the consecutive goals reached (*right*) shows the performance of the agent after training. Red lines indicate the median, white circles indicate means, and red triangles modes.

Note that Figure 7 shows two metrics of performance: The cumulative reward based on the task set in the example segment in Section 3.3, and the number of consecutive goals reached (CGR) within one episode. Although the former differs from the latter by incorporating the distance between initial position and goal, the two metrics provide nearly identical insights. Nevertheless, when communicating the performance of goal-driven models trained by RL, we recommend to report interpretable auxiliary metrics (like CGR for IHOM), instead of cumulative rewards. This abstracts away the actual performance from the specific reward function used and even allows one to compare their variants. Most importantly, however, it enables readers who are less familiar with reinforcement learning terminology or who may not be acquainted with the specific reward function to quickly assess the power of a model.

After training, we evaluated the best performing version of the agent (i.e., the set of weights that achieve the maximum average results in its gathering cycle) on 480 independent episodes. Here, actions are no longer sampled from the policy distribution, but instead the most probable action is chosen deterministically. The distribution of these episodes as measured by CGR is shown in Figure 7. It can be observed that while on average the agents chain approximately 34 reorientations, in most episodes they actually chain the maximum possible 50 reorientations, as discerned from the mode. The best agent is highly stable, with both the median and the mode at 50.

Our results show that while most agents converge reliably to a reasonable performance (30 CGR or higher), outliers can occur. Specifically, both time-to-convergence and peak performance vary between runs. Designing plausible models can mitigate this variance. By constraining the flow of information and its integration, the policy space can be transformed to favor specific solutions to the problem. At the same time, as policies are similar in their manipulation strategy but only differ in the effectiveness of their execution, the observed variance between agents can be interpreted as an inter-individual difference in a neuroscientific context.

The architecture we employed in this example was proposed in the seminal work of OpenAI et al. (2020). It is *not* biologically plausible but serves as a good baseline for assessing our tool. The training performance that we report in Figure 7 differs from OpenAI et al. (2020) primarily because the network relies only on raw sensory information and, second, because batch sizes and the amount of data gathered per cycle differ for practical, hardware-related reasons.

### 4.2 Benchmarks

AngoraPy aims to provide out-of-the-box functionality on a wide range of tasks. The following section presents benchmarking experiments on classical and sensorimotor tasks to demonstrate the general applicability of AngoraPy to various problems beyond the one showcased above. Naturally, this set of tasks is not exhaustive. Nevertheless, the following indicates the general-purpose applicability of the stack of methods underlying AngoraPy’s training framework.

#### Classical Control

Pendulum equilibration tasks constitute a standard benchmark for control systems. We present learning trajectories for the Gym implementation of three variants in Figure 8 alongside a classic trajectory optimization for a 2D spacecraft (LunarLander). Without any specific parameter tuning, AngoraPy’s PPO implementation solves all tasks.

**Figure 8.**
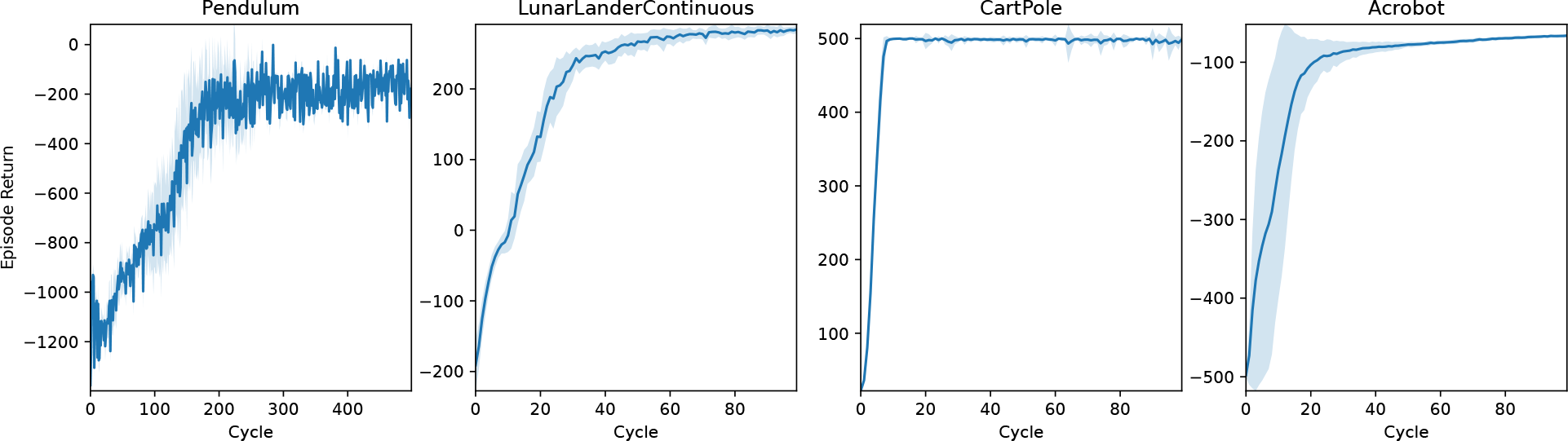
Learning curves of agents trained on different classical control tasks. All agents use the same two-layer neural network architecture with a Beta/categorical policy head. Weights were not shared between policy and value network. Blue lines represent the average returns of 16 independent agents, and the lighter area around them is their standard deviation.

#### 3D Control in MuJoCo

Whereas the control tasks in Figure 8 are easy to solve for most state-of-the-art RL algorithms, the set of (robotic) control tasks benchmarked in the first three columns of Figure 9 pose a more significant challenge. This challenge stems i) from a more rigid simulation in three-dimensional space, ii) from purely continuous control, and iii) in some cases, from higher degrees of freedom. Nevertheless, PPO again demonstrates an impressive invariance in its applicability to different robots using the same parameters. Note that for most of the environments tackled in Figure 9, the original paper introducing PPO (Schulman et al., 2017) already presented benchmarks, yielding similar results with differences stemming from different parameter settings. Agents using Beta distributions often outperform or are on par with their Gaussian counterparts on these tasks.

**Figure 9.**
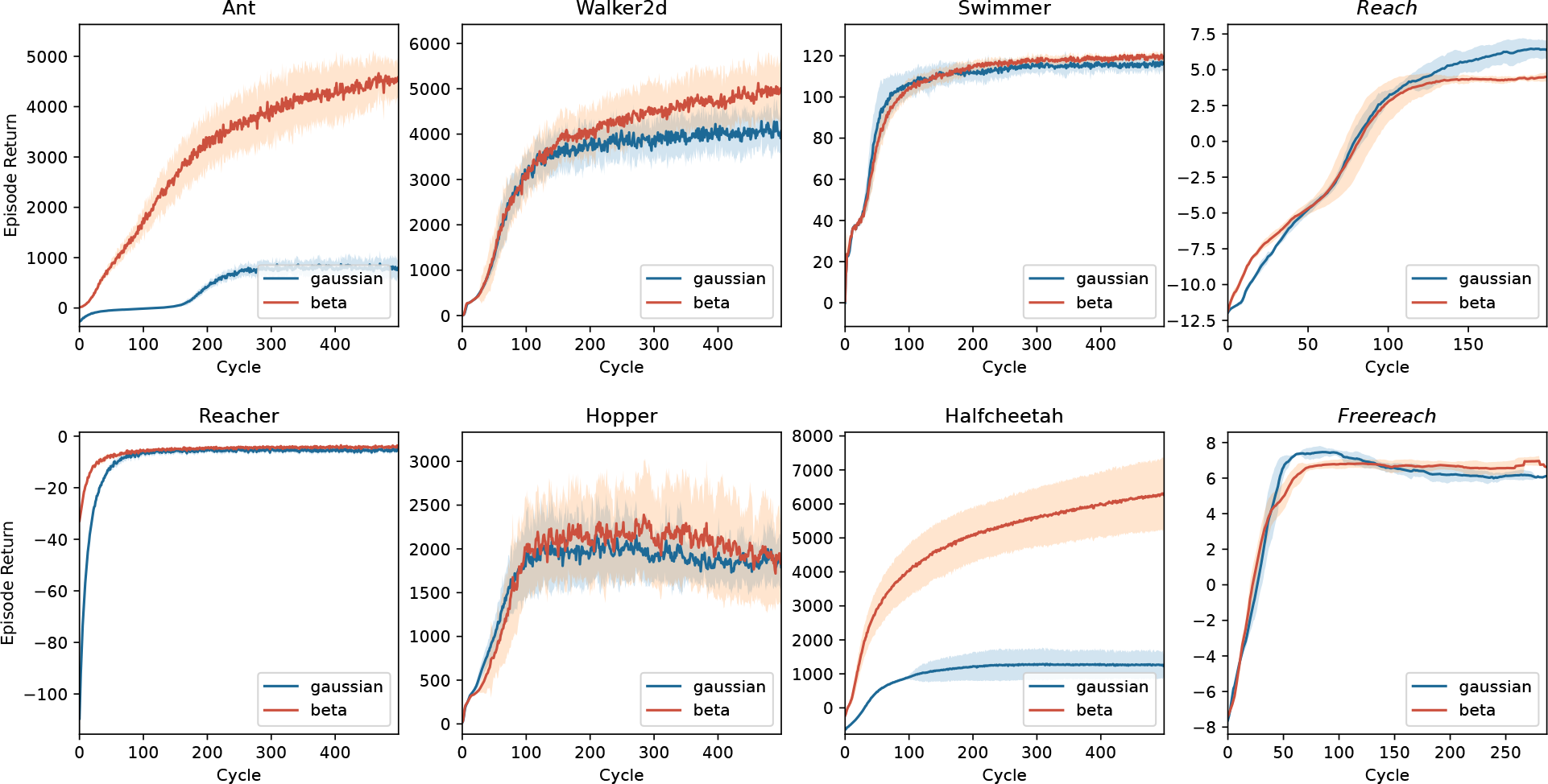
Benchmark experiments on three-dimensional control tasks simulated in MuJoCo. Tasks with *italic* titles are anthropomorphic reaching tasks. In all other tasks, the model controls non-anthropomorphic motor plants. The latter agents use the same two-layer architecture used in Figure 8. For anthropomorphic tasks, a recurrent network with an LSTM cell builds the policy. Lines represent the average returns of 16 independent agents, and the lighter area around them is their standard deviation. Blue lines correspond to agents using a Gaussian policy, whereas red lines summarize the performance of Beta distributed policies.

#### 20-DoF Dexterous Control

One of the most fascinating and complex problems in the motor domain is sensation-guided manual control. The complexity of the hand as motor plant stems from high degrees of freedom, coupled with non-trivial dependencies between joints. To solve any task involving this plant, the model first needs to learn the relationship between control applied to the joints and its effect on the position of the plant’s individual parts. A task focusing on this raw skill thus is a good benchmark and initial proof-of-concept for any model of the sensorimotor system. In Figure 9, the last column shows results for two manual dexterity tasks, *reaching* and *freereaching*. Both tasks challenge the agent to join the thumb’s fingertip with one other digit’s fingertip (target finger) as indicated by the goal description. In *reaching*, we model this task by rewarding the proximity of individual fingertips to locations sampled around their initial position for non-target fingertips and sampled around a meetup point for the thumb’s and the target’s fingertip. In *freereaching*, we abstract away from predetermined locations and instead formulate the reward as the inverse distance between the thumb and the target finger, and add a punishment for other fingertips entering a zone around the thumb’s fingertip. As demonstrated in Figure 9, both formulations of the task can be reliably learned within 100 *−* 150 cycles.

## 5 Discussion

We introduced *AngoraPy*, a Python library for building goal-driven models of the human sensorimotor system at scale. AngoraPy provides a distributable reinforcement learning backend accessible through a high-level API. It thereby allows training of large recurrent convolutional neural networks (RCNNs) as models of the brain and features standards for defining model architectures and tasks. Tasks are implemented on top of Gym (Brockman et al., 2016), and bodies and their environments are simulated in MuJoCo (Todorov et al., 2012). A synergy of wrappers and interfaces standardizes the setup of anthropomorphic task environments. These ship together with predefined environments for dexterity tasks. Sensory modalities comprise vision, touch, and proprioception, or a subset thereof. Motor commands are issued by stochastic policies and sampled from multivariate distributions over the joint space of the motor plant. The construction of models is standardized but flexible. AngoraPy supports multi-modality, weight sharing, asymmetric policies, and pretrainable and preloadable components. The reinforcement learning backend implements proximal policy optimization (Schulman et al., 2017) and embeds it in a stack of supportive methods. To support large RCNNs, the gradient-based optimization chunks sequences and truncates out-of-chunk timesteps. Both simulation and optimization are highly scalable via native MPI distribution strategies. Taken together, AngoraPy combines the power of state-of-the-art reinforcement learning with high-performance computing in a sensorimotor modeling toolbox.

We have validated AngoraPy’s ability to train arbitrary network architectures on several task domains, including anthropomorphic robots with up to 20 degrees of freedom. Our results highlight that no extensive hyperparameter tuning is necessary to successfully apply AngoraPy to new tasks, making the toolbox particularly suitable for users who are not experienced with reinforcement learning. A case study on in-hand object manipulation detailed the use of AngoraPy for studying intricate anthropomorphic motor functions.

Previously, there was a lack of software supporting modeling in the sensorimotor domain. While several libraries exist that implement various RL algorithms (e.g., Raffin et al., 2021; Moritz et al., 2018; Guadarrama et al., 2018), these libraries are primarily intended for RL researchers. This is reflected in the wide range of algorithms offered, which can come at the cost of user-friendliness for non-experts. It additionally forfeits flexibility with respect to applications in favor of flexibility with respect to algorithms. In consequence, applying an algorithm to a specific problem can become cumbersome if the flexible standards of the library do not align well with the requirements of the problem. In neuroscientific modeling, where RL algorithm choice is less important, a tailormade tool is thus more efficient and easy-to-use. Moreover, these libraries do not always allow for training the complex networks desirable to model the sensorimotor system. For instance, the PPO implementation by Raffin et al. (2021) cannot be applied to RNNs. Lastly, being non-specific in nature, they require substantial engineering when applied to training anthropomorphic sensorimotor models. With AngoraPy, we offer the research community a comprehensive tool tailored specifically towards its application in neuroscience. In this domain, our library thus bests aforementioned libraries in functionality and ease-of-use.

With its comprehensive set of features, AngoraPy can be used by neuroscientists who seek to build models of sensori-motor systems that bridge across Marr‘s (1982) levels of description of neural information processing. Specifically, our framework addresses the algorithmic implementation of a sensorimotor function guided by both computational theory and biological constraints. Observing under which constraints biologically valid organization emerges (and under which it does not) may additionally support our understanding of the interplay between structure and function. The resulting goal-driven models can then be utilized to evaluate existing hypotheses as well as to generate novel predictions to guide both theoretical and empirical research. For example, AngoraPy may be used to develop models of the frontoparietal network with biologically inspired pathways and regions trained on in-hand object manipulation tasks in order to better understand the neurocomputations that underly human dexterity. These models may then generate new hypotheses about information processing that occurs within and between areas in both feedforward and feeback directions. Similarly, AngoraPy may be employed to study complex motor skills such as grasping, reaching, manipulation, balancing, or locomotion.

## Author Contributions

TW developed the software introduced in the manuscript. TW wrote the first draft of the manuscript. TW and MS contributed to manuscript revision. MS and RG secured funding for the project. TW, MS and RG read and approved the submitted version.

## Acknowledgments

We acknowledge the use of Fenix Infrastructure resources, which are partially funded from the European Union’s Horizon 2020 research and innovation programme through the ICEI project under the grant agreement No. 800858.

## Data Availability Statement

Software introduced in this paper is available at https://github.com/ccnmaastricht/angorapy and licensed under GNU GPL-3.0.

## Footnotes

1 https://github.com/ccnmaastricht/angorapy

2 https://github.com/weidler/angorapy-tutorials

3 https://www.shadowrobot.com/dexterous-hand-series/

4 The number of GPUs in this setup is a byproduct of how the HPC architecture that we use is set up and should not be taken as a guideline. Usually, a substantially lower number of GPU optimizers should suffice.

